# Biomechanics of cutting: sharpness, wear sensitivity, and the scaling of cutting forces in leaf-cutter ant mandibles

**DOI:** 10.1101/2023.05.10.540164

**Authors:** Frederik Püffel, O. K. Walthaus, Victor Kang, David Labonte

## Abstract

Herbivores large and small need to mechanically process plant tissue. Their ability to do so is determined by two forces: the maximum force they can generate, and the minimum force required to fracture the plant tissue. The ratio of these forces determines the required relative mechanical effort; how this ratio varies with animal size is challenging to predict. We measured the forces required to cut thin polymer sheets with mandibles from leaf-cutter ant workers which vary by more than one order of magnitude in body mass. Cutting forces were independent of mandible size, but differed by a factor of two between pristine and worn mandibles. Mandibular wear is thus likely a more important determinant of cutting force than mandible size. We rationalise this finding with a biomechanical analysis which suggests that pristine mandibles are ideally ‘sharp’ – cutting forces are close to a theoretical minimum, which is independent of tool size and shape, and instead solely depends on the geometric and mechanical properties of the cut tissue. The increase of cutting force due to mandibular wear may be particularly problematic for small ants, which generate lower absolute bite forces, and thus require a larger fraction of their maximum bite force to cut the same plant.

## Introduction

Plant-feeding occurs at vastly different scales, from large bulk-feeding mammals to tiny cell-ingesting leaf miners [1]. Despite these differences in scale, all herbivores share the same basic task: they need to mechanically process the plant tissue; if they cannot tear, masticate, cut, pierce or drill into the plant, they cannot feed on it. From a simple mechanical perspective, a necessary condition for plant-feeding is then given by the ratio of two key forces: the maximum force the animal can generate needs to exceed the minimum force required to fracture the plant tissue [1–3]. How do these forces change with animal size?

Based on a simple scaling argument, the maximum available force is expected to increase in proportion to a characteristic area, or with body mass to the power of two-thirds [4]. However, the scaling of the fracture force is difficult to predict, because it depends on plant-mechanical properties [5–9], the mode of fracture [3, 5], and on the geometry of the cutting, chewing or piercing ‘tool’ in question [10–15]. In absence of robust theoretical frameworks, fracture forces are often determined experimentally instead [e. g. 12, 14, 16, 17]. A key challenge for such experimental approaches is that studies across a large tool size range typically require using different species, so that fracture tools usually differ in both scale and shape [2, 12]. In order to investigate the influence of tool size alone, we here measured fracture forces using the cutting tools of a species for which adults vary substantially in size but only little in shape: *Atta vollenweideri* leaf-cutter ants.

Leaf-cutter ant colonies consist of up to several million workers, which cover a large range of body sizes, from less than a milligram to over 100 mg in some *Atta* species [18–21]. Notably, this size range reflects ‘static’ differences among workers of equivalent developmental stages; workers retain their adult form after eclosion from the pupa. Fully-matured leaf-cutter ant foragers cut leaf fragments from plants in the colony surroundings; these fragments are then carried back to the colony to feed them to a subterranean fungus grown as crop [22–25]. To cut transportable fragments from large leaves, leaf-cutter ants typically use one of their mandibles as an ‘anchor’ which pierces through the leaf lamina but remains approximately static; a single cut is then made by drawing the second mandible through the leaf lamina like a blade [26, 27, see SI video]. Repeated cutting cycles, combined with a ‘pivoting’ of the ant around an approximately fixed anchor point for the hind legs, then yields leaf fragments with semi-circular shape that can be carried back to the nest [e. g. 26, 28]. Interestingly, the tendency to cut and carry plant fragments correlates with worker size: larger ants cut and carry larger fragments [19, 28–30], at higher speeds [27, 30–35], and forage on ‘tougher’ plants than smaller ants [23, 24, 28, 31, 33, 36–38].

In contrast to this robust empirical evidence for size-related preferences in foraging, the biomechanical factors that underpin it remain poorly understood [but see 27, 28, 39]. For example, do larger workers cut tougher leaves because smaller workers are unable to do so, or because they are more efficient? In order to assess how the ability to cut leaves varies with size, we previously measured the maximum bite forces of *A. vollenweideri* leaf-cutter ants [40]. Peak bite forces increased with strong positive allometry, *F*_*b*_ ∝ *m*^0.90^, in substantial excess of the isometric prediction, *F*_*b*_ ∝ *m*^0.67^: A large forager of 40 mg generates peak bite forces of about 800 mN, 16 times more than a small forager of 2 mg, *F*_*b*_ ≈50 mN, and about as large as the bite forces of a vertebrate 20 times heavier [40]. As a result, large foragers are presumably able to cut a considerably larger fraction of tropical leaves [8, 40].

However, this conclusion is speculative, because it remains unclear how the forces required to cut vary with mandible size. For example, one may speculate that cutting forces vary with a characteristic length [e. g. 12, 41, 42]; smaller ants would then have ‘sharper’ mandibles which demand less force to cut a given material. To complicate matters further, mandible ‘sharpness’ may vary across the lifetime of an ant due to mandibular wear [27]. The degree of mandibular wear likely depends on the abrasiveness of the cut materials [43, 44], the wear resistance of the mandible teeth [45–47], the mandible tooth geometry [2, 12], and the forces involved in cutting [12]. To investigate the impact of mandibular wear on cutting forces and to compare it to the impact of size, we performed cutting force experiments us-ing either mandibles from freshly eclosed ants (callows), which initially remain in the nest and thus have ‘pristine’ mandibles, or from workers which actively partook in foraging, and thus are likely to have worn mandibles. We hypothesise (i) that mandibles of small ants cut with less force because they are sharper, and (ii) that forager mandibles require the application of larger forces compared to callow mandibles of the same size, as they are blunted by wear.

## Materials & methods

### Study animals

We sampled *A. vollenweideri* leaf-cutter ants from two colonies, founded and collected in Uruguay in 2014. The colonies were kept in a climate chamber (FitoClima 12.000 PH, Aralab, Rio de Mouro, Portugal) at 25 ^*°*^*C* and 50 - 60 % relative humidity, with a 12/12 h light-dark cycle. They were provided with bramble, laurel, maize and honey water *ad libitum*, supplied in a foraging arena that was connected to the main colony via PVC tubes (*≈*30 cm to the closest fungus box; 25 mm inner tube diameter).

To quantify the force required to cut thin leaf-like sheets with mandibles of different sizes, we collected two sets of ants across the worker size-range excluding the smallest workers, which typically do not cut leaves [body mass *<*1 mg, see 24, 48].

First, we extracted workers from the fungal garden that had either freshly eclosed, identified by their bright cuticle, or were still in the pupal stage [n = 46, 27, 49]. In the weeks following eclosion, callows remain inside the nest and abstain from foraging activities [24, 50]. The mandibles of callow workers are thus likely ‘pristine’, which allowed us to test for the effect of mandible size on cutting force without potentially confounding effects due to mandibular wear [27]. To ensure that the incorporation of zinc into the mandibular teeth was completed, callows were kept alive for at least 72 h post eclosion, defined as time point at which the legs had completely unfolded [27, 49]. To monitor pupae and callows, they were placed in centrifuge tubes, which in turn were kept inside the foraging arena. The tubes contained small amounts of fungus, and had a 3D-printed polylactic acid (PLA) lid with holes too small for the collected workers to pass through, but large enough for minims to enter for pupal maintenance [27]. This method was thus unsuitable for smaller ants (*<*10 mg), which were collected by transferring late-stage pupae into a separated container with sufficient amounts of fungus and numerous minims instead. Hatched ants were marked with a unique colour code [Edding 4000 paint marker, Edding AG, Ahrensburg, Germany; 51].

Second, we collected fully-matured workers from the foraging arena (n = 39). Depending on their age, these workers may have mandibles worn from the repeated cutting of leaves [27]. Quantifying the mandibular cutting forces for active workers allowed us to investigate the effect of mandibular wear and its interaction with worker size.

### Mandible preparation and wear quantification

All ants were sacrificed by freezing, weighed to the nearest 0.1 mg (Explorer Analytical EX124, max. 120 g x 0.1 mg, OHAUS Corp., Parsippany, NJ, USA; body mass ranged between 1.8 to 46.4 mg), and decapitated using micro-scissors. The head capsules were split in half along the sagittal plane using a scalpel, and only the left head hemisphere was retained (see Fig. 1A). Leaf-cutter ants show no preference between left and right mandible when cutting [26], and their bite apparatus is bilaterally symmetric [52]. We hence assume that there are no systematic differences between both sides. To facilitate sample mounting, insect pins were inserted into the head halves (size ‘2’ for ants *<* 10 mg, size ‘4’ for ants of 10-20 mg, and size ‘6’ for ants *>* 20 mg; Shigakontyu, Tokyo, Japan). The interface between insect pin, head capsule and mandible base was then immobilised with two-component epoxy to minimise compliance of the mandible-head-pin complex (Araldite Rapid, Huntsman Corp., The Woodlands, TX, USA; see Fig. 1A).

**Figure 1.**
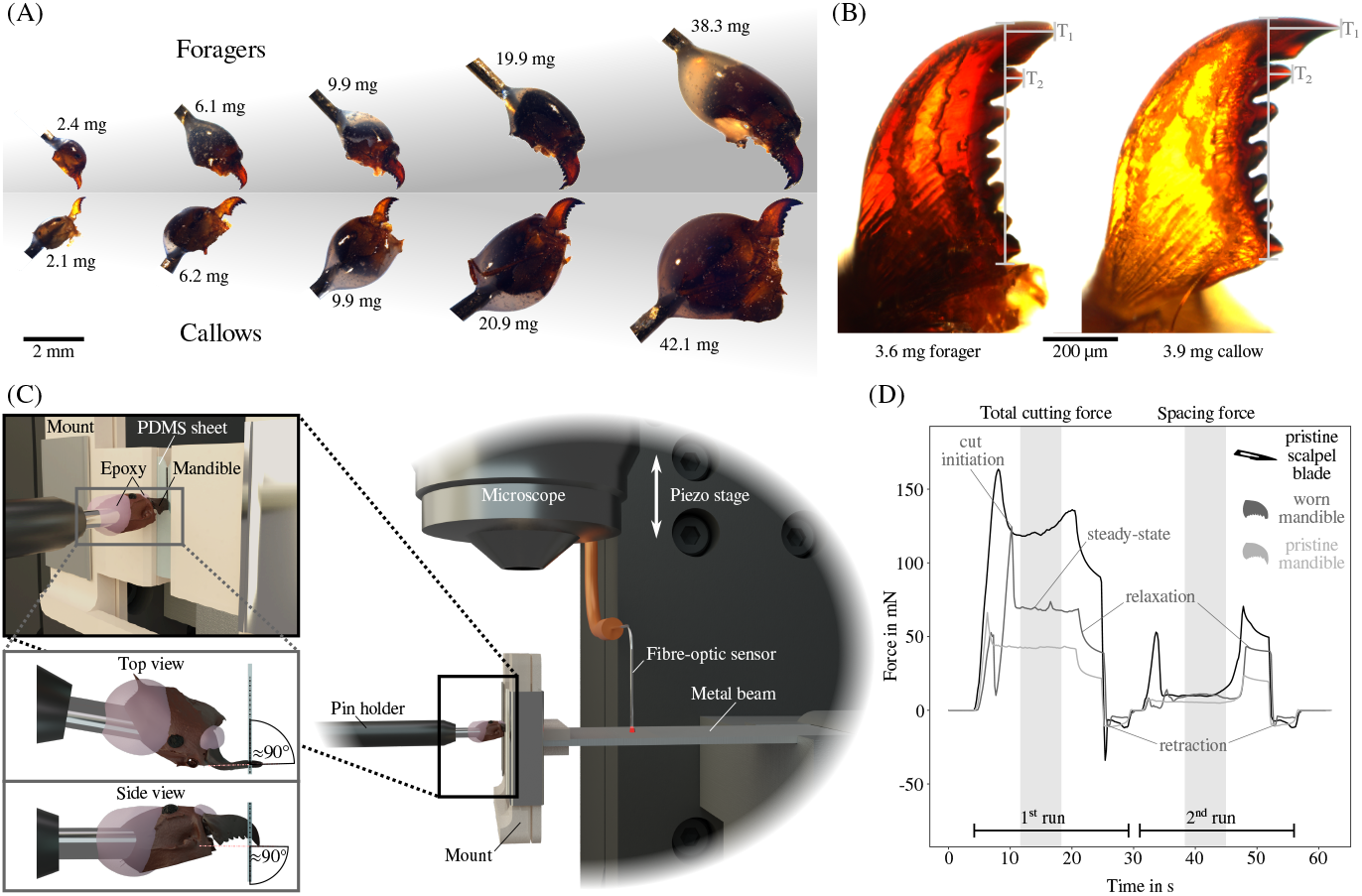
**(A)** In order to measure mandibular cutting forces, *Atta vollenweideri* leaf-cutter ants were extracted from the foraging arena and fungal garden of mature colonies (body mass: 1.8-46.4 mg). Both foragers (with worn mandibles) and callows (with pristine mandibles) were used for the experiments to quantify the effects of both worker size and mandibular wear on cutting force. **(B)** For each mandible, we calculated a wear index based on absolute length changes to the most and second most distal teeth, *T*_1_ and *T*_2_, respectively [see 27, and SI for more details]. **(C)** Cutting forces were then measured using a custom-built setup based on a fibre-optic displacement sensor and a bending beam, both connected to a piezo motor stage. A PDMS sheet was fixed in a custom-designed holder, mounted at the free end of the beam, and the mandible was positioned above the sheet such that its cutting edge was perpendicular to the sheet plane. The motor then moved the beam mounting vertically against the mandible, causing the sheet to be cut and the beam to deflect. **(D)** After an initial loading phase, cutting force peaked at cut initiation, and then dropped to an approximately constant value. At the end of this ‘steady-state’ phase, the forces dropped again when the motor stopped and became negative as the setup was moved back to its original position. A second run through the cut was performed in the same position to extract the spacing force ‘lost’ to sheet bending and friction [e. g. 11, 42]. The drift-corrected average total cutting force and the corresponding spacing force across 2 mm cutting distance (shaded areas) were extracted for further analysis

In order to determine a proxy for the degree of mandibular wear, all mandibles were photographed with a camera mounted onto a light microscope, such that their dorsal surface was in focus (DMC5400 on Z6 Apo, Leica Microsystems GmbH, Wetzlar, Germany; see Fig. 1B). Numerous empirical metrics for mandibular wear have been proposed in literature, including variation of mandible length [53, 54], shape changes of the mandibular cutting edge [55–57], number of lost mandibular teeth [58], reduction in profile area of distal mandibular teeth [59], and length changes of the mandibular teeth most relevant for cutting [27, 60]. All of these metrics are proxies with no direct established mechanistic relation to cutting force. As such, their predictive value can only be assessed in correlation to direct cutting force measurements, and selecting any one of them is difficult to justify *a priori*. We chose a metric that has been demonstrated to correlate significantly with cutting force in closely-related *A. cephalotes* ants, so enabling a direct comparison [27]; however, we do not intend to imply that this metric is more or less predictive than any of the others. Following Schofield et al. [27], a mandibular wear index, *W*, was thus defined as:

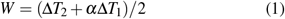

Here, Δ*T*_1_ and Δ*T*_2_ are the differences between observed and pristine tooth length for the most distal and second most distal tooth, respectively; *α* is a weighting factor, defined as ratio between the average length differences, 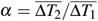 [for more details, 27]. This wear index has dimension length, and may be interpreted as the weighted average length change of the two distal-most teeth. This wear index definition thus is a proxy for absolute rather than relative wear.

To calculate the wear index, the length of the mandible blade and the two most distal teeth were measured from each photograph (see Fig. 1B), and the pristine tooth length as function of body mass was estimated via regression analysis on measurements of callow mandibles (for exact methodology, see SI). The wear index could not be extracted for 17 out of 76 mandibles, because relevant parts of the mandible were obstructed by the head capsule.

### Cutting force setup

Mandibular cutting forces were measured with a custom-built setup based on a fibre-optic displacement sensor (*µ*DMS-RC32 controlled via DMS Control v 3.015, Philtec Inc. Annapolis,MD, USA; linear range of 2.5 mm, recording at 81.4 Hz at 30^*°*^ and 50 % transmitted optical power). The sensor was held in place by a custom-built holder, mounted on two micromanipulators to control its orientation, and attached to a piezo motor stage (M-404.6PD controlled via PIMikroMove v 2.33.3.0, Physik In-strumente GmbH & Co. KG, Karlsruhe, Germany; see Fig. 1C). The sensor was placed above a stainless steal bending beam with a thickness of 0.35 mm, a width of 10.4 mm, and a free length of 28.7 mm, such that the sensor tip was about 400 *µ*m above the beam surface [see 61, for a similar setup]. The beam was clamped to the motor stage at one side. At the free end of the beam, a 3D printed mount was attached. This mount held the cutting substrate during the experiments, clamped in place by two metal clips on either side (Supaclip 40, Rapesco Office Products PLC, Sevenoaks, UK). At the centre of the mount, there was a ‘free’ cutting region (1.5 mm wide and 8 mm long), over which mandibles or scalpel blades were positioned for cutting experiments (see Fig. 1C).

The sensor was calibrated with a series of ten calibration weights ranging between 10-245 mN (1-25 g, Kern & Sohn GmbH, Balingen, Germany), covering the range of observed total cutting forces (19 - 172 mN). Weights were suspended from the mount in increasing order, and at the lever arm at which cutting forces were applied. For each calibration weight, we averaged the sensor output across 5 s after initial force fluctu-ations had faded (see SI Fig. 2C). To account for sensor drift, the sensor output was extracted for the unloaded beam at the beginning and end of the measurement, and a linear drift correction was implemented; sensor drift, however, was generally small, *≈* 0.01 mN s^-1^, or 6 mN min^-1^, and thus less than 5 % of the smallest total cutting force over the duration of a typical measurement of 60 s. From simple beam theory, the relationship between applied force and beam deflection should be linear for small deflections. We indeed observed linearity for calibration forces *<* 150 mN; for forces exceeding 150 mN, however, the sensor distance was systematically ‘sub-linear’, suggesting deflections sufficiently large to invalidate the use of the small angle approximation. We thus used a quadratic regression model to characterise the relationship between force and distance, which accounted for more than 99 % of the variation, and yielded a lower Akaike Information Criterion compared to a linear or cubic model (AIC_*linear*_ = 91.7, AIC_*quadratic*_ = 59.6, AIC_*cubic*_ = 61.6; see SI Fig. 2D).

### Polymer sheet production and mechanical testing

Previous studies on mandibular cutting forces used leaf lamina and floral petals as cutting substrates [26, 27]. This choice has the advantage that it is of direct biological relevance. However, plant tissues are typically heterogeneous, of uneven thickness, and have mechanical properties that vary with hydration and tissue age, so introducing potential for substantial covariation that is difficult to control [e. g. 9, 62–65]. In order to minimise variation due to material inhomogeneities, we used well-defined PDMS sheets as cutting substrate.

PDMS sheets were made with a 4:1 (silicon base: curing agent) mixing ratio (SYLGARD 184, Dow Inc., Midland, MI, USA). The mixed but uncured PDMS was sandwiched between two silanised glass plates, separated by feeler gauges (200*µ*m,Precision Brand, Downers Grove, IL, USA; see SI Fig. 2A), and pre-cured in an oven at 100^*°*^C for two hours (Drying oven, Sanyo Electric Co., Ltd., Osaka, Japan). The PDMS ‘sandwich’was then cooled to room temperature, slowly peeled from the glass plates, placed on aluminium foil, and fully cured at 165^*°*^C for a further 48 hours [66]. Sheet thickness was verified through measurement at six random locations across the sheet with a digital micrometer (max. 25 mm x 0.001 mm, Mitutoyo Corp., Kawasaki, Japan), and was 215± 8 *µ*m (mean±standard deviation), or within 10 % of the target thickness.

To mechanically characterise the PDMS sheets, pure shear tearing and uniaxial tension tests were conducted with a universal tension and compression system (Multitest5-xt, Mecmesin Ltd., Slinfold, UK; 10 N load cell and Mec277 double-action vice grips with diamond jaws). Two rectangular samples from each of the eight PDMS sheets were cut and used for pure shear tearing; in one of the two paired samples, a notch of 3 mm length was introduced at the centre of the short side (see SI Fig. 2A for dimensions). Both samples were tested at a small strain rate of 0.0067 s^*−*1^ (test speed divided by sample height) to approximate quasi-static loading conditions. The critical displacement to rupture was then extracted from the notched sample based on a time-synchronised video recording. The force-distance curve of the unnotched sample was integrated from zero to this critical displacement to obtain the work done by the applied load. Fracture toughness was then calculated as this work divided by sample width and thickness [for more details, see 67, 68], yielding an average of *G*_*c*_ = 98 ± 7 J m^-2^ [for comparison, see 68].

Next, uniaxial tension tests at 0.5 mm s^-1^ motor speed were conducted with two ‘dog-bone’ samples cut from each of eight sheets according to ISO standards (ISO37 and ISO5893). The Young’s modulus was extracted from the loading region of the stress-strain curve via linear regression between 0-10 % strain [69, 70]; on average, the Young’s modulus was *E* = 4.1± 0.3 MPa [in agreement with published values, 66].

### Cutting experiments

Individual ant heads were fixed onto a pin holder, which was connected to a 3D micromanipulator (n = 85; Manipulator MM 33, Märzhäuser Wetzlar GmbH & Co. KG, Wetzlar, Germany). The mandibles were then positioned using a top-down microscope such that the dorso-ventral head axis was approximately horizontal, the mandibular teeth were roughly perpendicular to the PDMS sheet, and the most distal tooth tip just about extended over the sheet edge [see 27, and Fig. 1C].

We cut a small wedge into all PDMS sheets (*≈* 30^*°*^ and 1.5 mm deep) to facilitate cut initiation by reducing effects of sheet bending and buckling [27, 68, 71]. The unstretched sheets were then placed individually between the two components of the polymer mount, and metal clips were slid onto the mount using the clip dispenser provided by the manufacturer, such that both clamps were approximately parallel and 6 mm away from the mount centre (see Fig. 1C); this procedure ensured that the clamping conditions were kept approximately constant across measurements.

The beam mount was then moved toward the mandible until the tip of the pre-cut wedge was about to contact the mandibular cutting edge. The sensor recording was started, and the beam mount was moved vertically against the mandible blade, resulting in cutting motion somewhat akin to the ‘blade-like’ cutting behaviour observed in freely cutting leaf-cutter ants [26, 27]. The motor moved at a constant speed of 0.3 mm s^-1^, at the upper end of cutting speeds observed during foraging [*≈* 0.02 - 0.30 mm s^-1^, 23, 26, 27, 30, 33, 34, 72], and over a total distance of 5 mm; the beam deflected by around 100 *µ*m for a medium cutting force of 65 mN, so that the corresponding displacement of the sheet-holding mount was about 4.9 mm (see SI Fig. 2C & D). The sheet was subsequently retracted to its original position, and a second run was initiated in order to extract the force due to elastic sheet deformation and sidewall friction [henceforth referred to as spacing force, e. g. 11, 42, 73, 74]. After a force peak at cut initiation, the total cutting force remained approximately constant until the motor stopped, and force decreased (see Fig. 1D). We extracted the drift-corrected steadystate total cutting force averaged across 2 mm following the initial peak; the corresponding spacing forces were extracted from the second run at the same motor positions, and averaged across the same distance (see Fig. 1D).

Cutting speeds typically vary with forager size; larger ants cut more quickly than smaller ants [27, 33, 34]. The effects of speed on cutting force depend on the viscoelastic properties of the material, but are typically small for elastomers such as PDMS cut at low rates [68, 75]. To briefly confirm that the speed-dependency is indeed small, we performed a series of measurements with the mandible of a single forager with a body mass of 19.9 mg at 0.1 mm s^-1^, 0.2 mm s^-1^ and 0.3 mm s^-1^ motor speed. Three repetitions were completed per speed, without remounting the mandible between measurements to reduce confounding effects due to small variations in mandible blade orientation. Variation due to remounting was quantified by measuring cutting forces of one small (5.4 mg) and one large forager (38.4 mg) at a constant cutting speed of 0.3 mm s^-1^. Both samples were mounted three times onto the pin holder, and cutting experiments were performed three times per mount.

Mounting had no significant effect on total cutting force (Analysis of Variance (ANOVA), small worker: F_2,6_ = 4.43, p = 0.07; ANOVA, large worker: F_2,6_ = 0.26, p = 0.71); we hence pooled the nine measurements per mandible and calculated the coefficients of variation, *CV*_small_ = 0.10 and *CV*_large_ = 0.03 (see SI Fig. 1). The relative force variation was significantly larger for the smaller mandible [Asymptotic test for equality of *CV* : D_*AD*_ = 7.76, p < 0.01, implemented in the R package ‘cvequality’, v 0.2.0, 76], suggesting that consistent mandible alignment is easier for larger mandibles. However, even for the smaller mandible, the force variation was small in comparison to the inter-individual variation across all foragers, *CV*_foragers_ = 0.52 (see below). We thus performed only a single measurement per specimen, unless otherwise indicated.

To contextualise our results based on *biological* ant mandibles and *synthetic* PDMS sheets, we performed two additional experiments. First, we measured cutting forces of pristine scalpel blades (Carbon steel, No.11, Swann-Morton Ltd., Sheffield, UK), positioned such that the blade tip just about extended over the sheet edge to reduce the contact area with the PDMS sheet (n = 5). Second, we performed cutting experiments with mandibles on a biological substrate, the leaf lamina of Japanese laurel, *Aucuba japonica*; the colonies were regularly fed with these leaves, and the lamina appeared comparatively homogeneous. Laurel leaves were cut from the plant on the day of the experiment, and kept hydrated using wet tissues between collection and measurement. To reduce variation due to material inhomogeneities, we cut all laurel samples from the same plant, from a leaf region close to the mid-vein. Prior to the cutting experiment, we measured lamina thickness and mounted the samples such that the cut ran perpendicular to the mid-vein. We used mandibles of 13 out of the 85 prepared ants, seven foragers (body mass 5.4 - 38.8 mg) and six callows (body mass 6.2 - 46.4 mg), mounted once with 1-3 repetitions per specimen. To account for differences in lamina thickness, *t*_*l*_, we corrected the measured total cutting force, 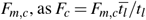, where 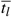 was the average lamina thickness (256 ± 29*µ*m).

Across all experiments, measurements were considered invalid and thus repeated when at least one of the following criteria was met: (i) the head capsule came into contact with the clamp or the cutting substrates; this occurred when the head capsule was initially close to the clamp and the PDMS sheet buckled; (ii) the mandible slipped out of the cut; (iii) the steadystate phase was too short to extract a meaningful cutting force (*<* 2 mm); (iv) the epoxy fixation of the joint failed, leading to mandible rotation; in these cases, the samples were re-glued and used again; and (v) the sample slipped out of the pin holder as observed occasionally for measurements involving high cutting force as on laurel (see below).

### Data curation and statistical analysis

We excluded a total of four out of 46 callows and five out of 39 foragers, because optical inspection of the mandible suggested that small amounts of epoxy contaminated the mandibular teeth, and cleaning attempts failed or caused visible damage. Additionally, we excluded one out of seven forager-laurel measurements, because the total cutting force exceeded the calibration range (*>* 245 mN).

Extraction of the average total cutting and spacing force from the raw data was done in python [v 3.9.7, 77], and all statistical analyses were conducted in R [v 4.1.1, 78]. To characterise the relationship between the extracted forces, body mass, and the two experimental groups (foragers vs callows), we used Analysis of Covariance (ANCOVA) with Type III sums of squares [79]. In addition, we performed Ordinary Least Squares (OLS) regressions to characterise the scaling relationships within the experimental groups. Unless stated otherwise, we performed these analyses on log_10_-transformed data.

## Results

### Cutting forces are independent of mandible size

Total cutting forces, *F*_*c*_, were independent of body mass (ANCOVA: F_1,72_ = 0.97, p = 0.33), but depended significantly on the experimental group (callows, vs forager, *F*_1,72_ = 21.2, p < 0.001; see Fig. 2A). These main effects must be interpreted with caution as the interaction term was significant [F_1,72_ = 4.42, p < 0.05, see 79], suggesting that the relationship between total cutting force and body mass may differ between the experimental groups. Indeed, within callows, total cutting forces tended to increase with body mass, whereas they decreased slightly within foragers. However, neither result was significant (p≥ 0.11, see Table 1).

**Figure 2.**
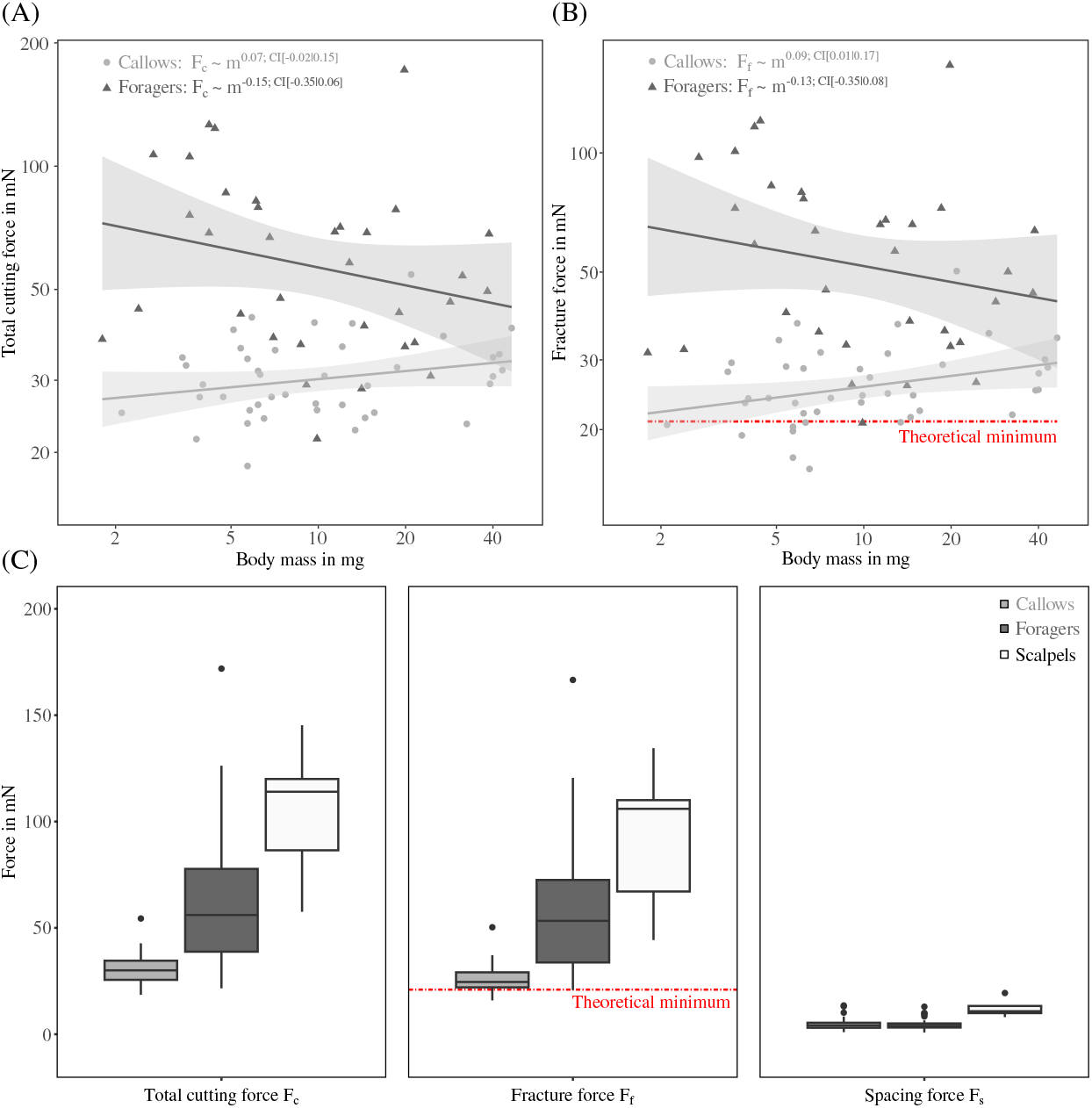
**(A)** Leaf-cutting is performed by workers spanning approximately one order of magnitude in body mass. To assess how cutting ability is affected by body mass, we measured mandibular cutting forces across almost the entire size-range (body mass, *m*, 1.8 to 46.4 mg), and across two experimental groups: callows with pristine mandibles (n = 42), and active foragers with mandibles affected to varying degrees by wear (n = 34). Total cutting forces, *F*_*c*_, were independent of body mass for both experimental groups (see main text for statistics), but twice as high for foragers compared to callows. **(B)** Fracture forces, *F*_*f*_, were not significantly affected by body mass in foragers. For callows, however, *F*_*f*_ increased significantly, *F*_*f*_ ∝ *m*^0.09^, from values close to a theoretical minimum based on pure shear tearing tests to values closer to those obtained from foragers. **(C)** On average, total cutting and fracture forces of both groups were significantly smaller than those measured for pristine scalpel blades (n = 5, *F*_*c*_ = 105±34 mN, *F*_*f*_ = 92±36 mN). Spacing forces, *F*_*s*_, were about 5±3 mN for both groups independent of body mass, and significantly smaller than for scalpel blades. Spacing forces contributed around 10 % of the total cutting force for mandibles.

**Table 1.**
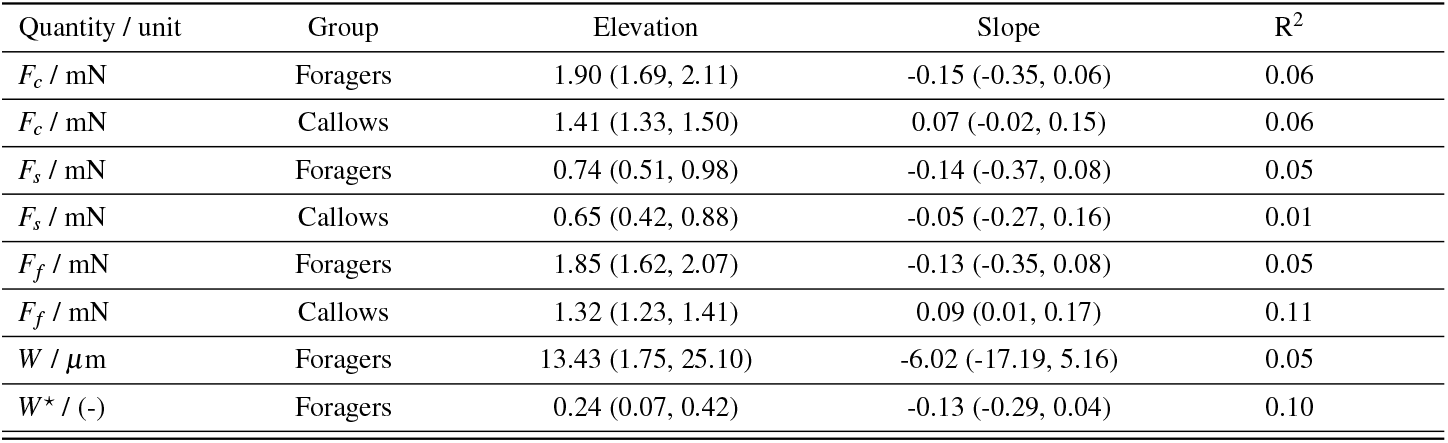
Results of Ordinary Least Squares regressions describing the relationship between total cutting force, *F*_*c*_, spacing force, *F*_*s*_, frac-ture force, *F*_*f*_, absolute and relative mandibular wear index, *W* and *W*^⋆^, respectively, with body mass in mg. All regressions were per-formed on log_10_-transformed values, apart from mandibular wear which contained negative values; this regression was done on semi-log_10_-transformed data instead. 95 % confidence intervals are provided in parentheses. The low R^2^ values underline that body size only had a small influence on all performance metrics.

Averaged among experimental groups, total cutting forces of foragers exceeded those of callows by a factor of two, 64±33 mN vs 31 ± 7 mN, respectively (see Fig. 2C). Notably, the coefficient of variation also differed significantly by about a factor of two (*CV*_foragers_ = 0.52 and *CV*_callows_ = 0.23; Asymptotic test for equality of *CV* : D_*AD*_ = 19.1, p < 0.001), suggesting that both relative and absolute force variation was larger among foragers. Despite these differences, the magnitude of total cutting force extracted for both groups was small in comparison to that of pristine scalpel blades, 105±34 mN, (see Fig. 2C; Wilcoxon rank sum test, forager mandibles vs scalpel blades: W = 29, p < 0.05; callow mandibles vs scalpel blades: W = 0, p < 0.001).

Across all ant mandibles, spacing forces, *F*_*s*_, were 5±3 mN with neither significant differences between experimental groups, nor significant size-effects (ANCOVA, experimental group: F_1,72_ = 0.34, p = 0.56; body mass: F_1,72_ = 0.25, p = 0.62; see Table 1 and SI figure). Because callow mandibles cut with less force, the relative spacing component was about two times higher (15 ± 8 % vs 8 ± 6 % (ANCOVA: F_1,72_ = 5.98, p < 0.05). Spacing forces of scalpel blades were 12 ± 4 mN, or 14 ± 8 % of the cutting forces, significantly larger than for callow and forager mandibles (see Fig. 2C; Wilcoxon rank sum test, forager mandibles vs scalpel blades: W = 8, p < 0.001; callow mandibles vs scalpel blades: W = 10, p < 0.001).

Because mandible spacing forces were size-invariant, the scaling of fracture forces, *F*_*f*_ = *F*_*c*_*−F*_*s*_, essentially mirrored the results obtained for the total cutting force (see Fig. 2B). Fracture forces were independent of body mass (ANCOVA: F_1,72_ = 1.64, p = 0.20), but depended significantly on experimental group (*F*_1,72_ = 22.6, p < 0.001), with a significant interaction (*F*_1,72_ = 4.46, p < 0.05). Within foragers, *F*_*f*_ tended to decrease with size, but this trend was not significant (p = 0.22, see Table 1); on average, *F*_*f*_ was 59±32 mN, comparable to the minimum force obtained for scalpel blades (44 mN). Within callows, however, *F*_*f*_ now increased significantly with worker size (p < 0.05, see Table 1), at the lower end approaching the minimum cutting forces predicted from tearing experiments (21 mN, see Fig. 2B and discussion).

To test if the observed differences in mandibular cutting force between callow and forager mandibles is also present with biological substrates, we measured cutting forces for a small subset from both experimental groups with laurel leaf lamina. Total cutting forces, corrected for differences in lamina thickness, were 141 ± 44 mN for foragers, exceeding those of callows (95 ± 14 mN) by almost 50 mN (see Fig. 3A); this difference was not significant (Welch Two Sample t-test: t_5.99_ = -2.40, p = 0.054). However, after subtracting spacing forces, the difference in fracture force was significant (114 ±23 mN vs 85 ± 11 mN; Two Sample t-test: t_10_ = -2.68, p < 0.05), with an average of 28 mN, similar to the result obtained for PDMS (33 mN).

**Figure 3.**
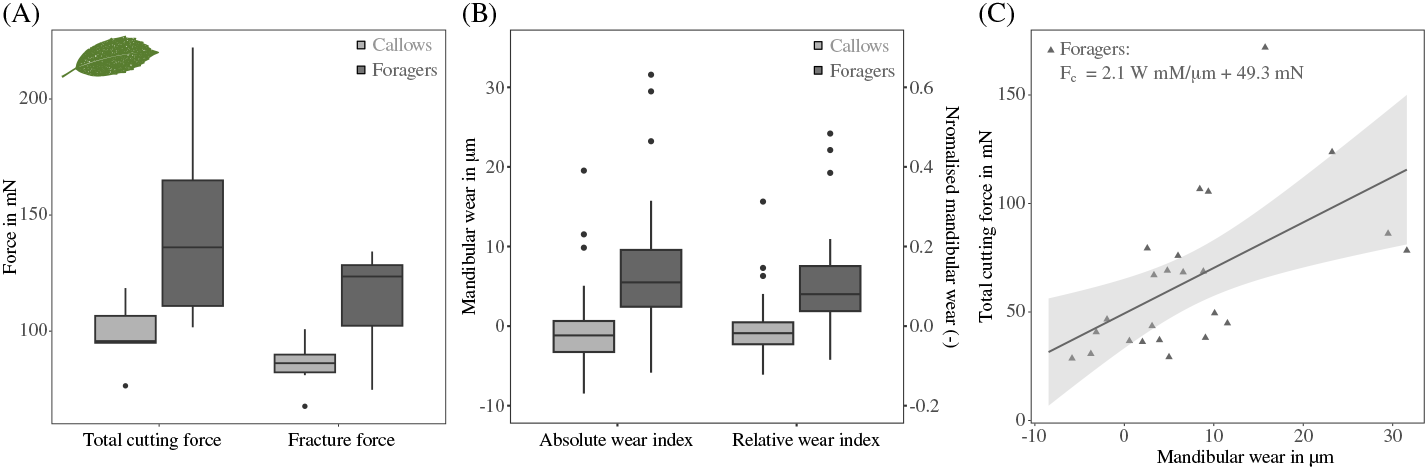
**(A)** We tested if the difference in cutting force between callows and foragers persists with biological substrates by performing cutting experiments on *Aucuba japonica* leaf lamina with mandibles from six foragers and six callows across the size-range. The absolute difference in fracture force between callows and foragers was 28 mN, similar to the results obtained for PDMS sheets (33 mN). **(B)** We calculated a simple mandibular wear index, defined as the weighted average length change of the two most distal teeth [n = 66, see 27, and Eq. 1]. In foragers, the absolute wear index, *W*, was 8 ± 10 *µ*m, independent of worker size; by definition, *W* was centred around zero for callows (0 ± 5 *µ*m). On average, foragers lost approximately 12% of their distal teeth in length, as indicated by the relative wear index, *W*^⋆^. **(C)** Mandibular cutting forces increased significantly with absolute wear index (n = 24, Ordinary Least Squares regression on forager data: slope = 2.09, 95 % CI (0.76s | 3.43), R^2^ = 0.33). Although the total variation explained by wear index remains below 50 %, it accounts for six times more variation than body mass.

### Cutting speed only has a small effect on cutting force

The cutting speeds during natural foraging typically vary with both worker size and leaf-mechanical properties; larger ants cut faster than smaller ants, and ‘tougher’ leaves are cut more slowly than ‘tender’ leaves [27, 33, 34]. We quantified the interaction between speed and total cutting force on synthetic PDMS sheets: Total cutting forces increased significantly but modestly with speed (ANOVA: F_1,7_ = 33.0, p < 0.001) from 29 ± 2 mN at 0.1 mm s^-1^ to 36 ± 1 mN at 0.3 mm s^-1^ (see SI Fig. 1). Total cutting forces thus increased by 20 % for a threefold increase in speed.

### Cutting forces increase significantly with mandibular wear

The mean mandibular wear index of foragers was 8±10 *µ*m, significantly different from zero, defined as the pristine state (One-sided Wilcoxon rank sum exact test: V = 272, p < 0.001), and independent of body mass (ANOVA on semi-log_10_-transformed data: F_1,22_ = 1.25, p = 0.28; see Table 1 and Fig. 3B). This size-independence suggests that absolute wear was the same across sizes, and thus that smaller ants lost a larger fraction of their teeth to wear. Although the relative mandibular wear index, normalised with the pristine length of the second most distal tooth, indeed slightly decreased with size from about 20 % for a 3 mg forager to 5 % for a 30 mg forager, this decrease was not significant (ANOVA on semi-log_10_-transformed data:F_1,22_ = 2.43, p = 0.13; see Table 1). Total cutting force increased significantly with absolute wear at a rate of 2.09 mN *µ*m^-1^ (OLS regression on untransformed data: 95 % CI of slope (0.76|3.43), p < 0.01, R^2^ = 0.33; see Fig. 3C), comparable to the rate of 3.7 mN *µ*m^-1^ reported for closely related *A. cephalotes* [27].

## Discussion

Leaf-cutter ants are iconic herbivores, with key impact on ecosystem ecology throughout the Neotropics [25, 80]. The continuous size-variation of their workers has also made them a model system for the study of ergonomic benefits of advanced polyethism in social insects [e. g. 24, 36, 81–83]. A key task faced by any leaf-cutter colony is to cut fragments in the colony surroundings, to maintain a fungus used as crop. Workers of which size are best suited for this task? Larger workers generate larger bite forces, and may thus be able to cut a larger variety of leaves [40, 52]. But the ability to cut depends not only on the available bite force, but also on the force required to cut the leaf with their mandibles – the key determinant is the ratio between both forces. Larger mandibles are putatively blunter, and may thus require larger bite forces to cut the same material [12–14]. How do cutting forces vary with mandible size?

In this work, we approached this question empirically, and measured the forces required to cut homogenous PDMS sheets with mandibles of workers of different body sizes. Cutting forces varied only weakly with mandible size, but differed considerably between mandibles taken from callows, which were pristine, and mandibles taken from foragers, which were affected to varying degree by wear. Before we discuss the biological implications and mechanical basis of these results, we briefly address two key aspects in which our experiments differ from natural cutting behaviour.

First, one may raise reasonable doubts about the extent to which results obtained on a synthetic elastomer can enable conclusions about biologically relevant cutting performance on leaves. The choice of PDMS as a test substrate was motivated by the need to minimise confounding variation in cutting forces due to material inhomogeneities, age- and hydration-dependence, expected for heterogeneous biological materials such as leaves [e. g. 8, 9, 62–65]. However, whether mandibles cut PDMS or leaves, the involved forces are amenable to mechanical analysis from first principles. We provide such an analysis at the end of the discussion, and the results confirm that the main conclusions of our study likely port to biological substrates, so enabling an initial discussion which focusses on biological implications.

Second, we acknowledge that even if experiments with PDMS can provide insights into cutting forces expected for leaves, our cutting experiments do not fully mirror the complexity of cutting behaviour of leaf-cutter ants. For example, mandibles rotate instead of translate; neck muscles may be used to change head and mandible orientation during cutting, and perhaps even directly contribute to cut propagation; and the section of the mandible blade used for cutting may be adjusted to account for local differences in mandible ‘sharpness’, or to dynamically alter the effective mechanical advantage of the mandible lever system. Despite these differences, two arguments suggest that our experiments are informative: Cutting forces of pristine mandibles were close to a theoretical minimum for PDMS; and although more complex mandible motion may decrease cutting forces in some cases [84, 85], out-of-plane forces applied to thin sheets likely result in sheet bending instead of concentrating tensile stresses, leading instead to an increase in cutting forces. Because leaf-cutter ants already need to show exceptional morphological and physiological adaptations to be able to produce bite forces sufficient to cut leaves [40, 86], it is biologically implausible and physically impossible that forces during ‘free cutting’ are substantially amplified over the minimum force dictated by leaf toughness (see below for a detailed quantitative argument).

### Size-invariance of cutting forces puts larger workers at an advantage

The weak size-dependence of cutting forces stands in stark contrast to the strong positive allometry of maximum bite forces in *A. vollenweideri*, which grow in almost direct proportion to body mass, *F*_*b*_ ∝ *m*^0.9^ [40]. As a result of this difference, the fraction of the maximum bite force required to cut the same material will decrease almost in direct proportion to mass, *F*_*c*_*/F*_*b*_ ∝ *m*^*−*0.9^ – a factor of 30^0.9^ *≈* 20 across the size range considered in this study. For materials that could in principle be cut by workers across the size range, the differential scaling of bite and cutting forces affords considerable behavioural flexibility to larger workers, bound by two extreme choices.

First, larger workers may choose to bite with maximum force, i.e. fully activate their closer muscles during cutting. The excess force *F*_*b*_*/F*_*c*_ directly determines the maximum possible strain rate of the mandible closer muscle during cutting [87]; larger ants would then cut with substantially larger speeds. Cutting speed amplification may be attenuated by viscoelastic effects which incur speed-dependent losses that increase the cutting force. In fracture, viscoelastic losses amplify the critical force by some power of the crack speed, and typically, 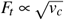 [88–90]. However, cutting forces usually show a much smaller speed-dependence, as the characteristic crack dimensions are tied to cutting tool geometry instead [e. g. 68, 75]. Indeed, a threefold increase in cutting speed resulted in an increase in cutting force of only 20 %, compared to about 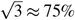 expected for tearing [68, and see SI Fig. 1].

Second, and alternatively, large workers may choose to bite with the same multiple of the required cutting force than small workers, i. e. only sub-maximally activate the mandible closer muscle during cutting, in which case muscle strain rate would be identical [87]. Although the cutting speed of larger ants would be sub-maximal as a result, this may be energetically advantageous, because muscle operates with maximum mechanical efficiency – the ratio between metabolic energy expended and mechanical energy produced – over a narrow range of intermediate strain rates [91]. On the basis of these arguments, we surmise that, even where a leaf can in principle be cut by small workers, it may be advantageous to assign larger workers to the task. In practise, foraging is a complex behaviour, and the behavioural choices of workers and their impact on the scaling of cutting speed and mechanical efficiency need to be addressed experimentally in future work.

### Cutting force variation is mainly driven by mandibular wear rather than body size

Throughout their life-time, leaf-cutter ant workers may cut a substantial amount of leaf tissue. To give a rough estimate, a mature colony of about one million foragers may cut about 3000 m^2^ of leaf area per year [25], and each square meter may require *≈* 3 km of cutting [92]. Across the time period where a worker may actively forage [about 4 months, 50], it may thus cut approximately 3 m leaf tissue, (4·3·3·10^6^)*/*(12·10^6^) = 3, or about 500 times their body length [*≈* 6 mm for a typical *A. vollenweideri* forager, 93]. Such extensive leaf-cutting likely causes substantial mandibular wear [27]. Consistent with this conjecture is the observation that average cutting forces of pristine and forager mandibles differed by about 35 mN, or a factor of about two for PDMS sheets, comparable to results on leaf lamina reported for closely related *A. cephalotes* [27]. The absolute difference may sound small, but it amounts to about 50% of the force required to cut the median tropical leaf, and to about 15% of the maximum bite force of a medium 10 mg forager [8, 40]. In absence of a strong size-effect, it appears that most of the difference between pristine and forager mandibles stems directly from mandible wear, or natural, wear-independent variation in mandible geometry. Indeed, even a simple empirical wear index based on the weighted average length change of the two distal-most teeth captures a remarkable 30% of the variation in cutting force, in striking contrast to the meagre 5-10% of variation explained by body mass (see see Table 1).

The substantial effect of wear on cutting forces is biologically meaningful, for it implies that wear may almost compete with body size in determining the ability of a worker to cut a given substrate: Cutting forces for mandibles from workers with a body mass between 4-6 mg varied by a factor of seven (n = 14), equivalent to the difference in maximum bite force between two workers that differ in mass by about a factor of 7^1*/*0.9^ ≈ 8 [40]. The effect of wear can thus be as large as the effect of an eight-fold reduction in the effective physiological cross-sectional area of the mandible closer muscle [52]. Both the susceptibility and the exposure to wear may themselves be size-dependent, putting smaller workers at further disadvantage. Mandibles of smaller workers may be more susceptible to wear, because they have to exert similar forces, but have smaller characteristic dimensions [12, 27]; they may be more likely to be worn, because foraging parties tend to be dominated by ants of intermediate size (between 3-10 mg, O.K. Walthaus et al., unpublished data). In support of this hypothesis, three lines of evidence may be presented: First, in large workers (body mass > 30 mg, n = 10), cutting forces varied only by a factor of three across pristine and forager mandibles, as opposed to a factor of seven for the mandibles of small workers (body mass < 6 mg, n = 24, see Fig. 2A). Second, although the scaling coefficients for total cutting forces of both callows and forager mandibles were not significantly different from zero (see Table 1), they were significantly different from each other (see results). Third, both absolute and relative wear index tended to decrease with size, although these trends were not significant (see results and **SI** figure).

Based on the significant increase of required cutting force with mandibular wear, we may speculate on the effect wear has on the fraction of cuttable leaves for both small and large workers. Previous analysis of leaf-mechanical data, in combination with bite force experiments, suggested that a 30 mg worker may be able to cut almost all species of tropical leaves, and a 3 mg worker may be able to cut about half of them [8, 40]. Although this analysis neglected the effects of friction and mandible geometry, it may still serve as a reasonable starting point to estimate the effects of wear. We may calculate the reduction in the fraction of cuttable leaves based on the following two assumptions: First, the required cutting force for a pristine mandible, *W* = 0, is size-invariant and approximately equal to the product between fracture toughness and leaf lamina thickness (also see below). Second, the increase in cutting force with wear is material-independent and equal to the regression slope extracted for PDMS (2.09 mN *µ*m^-1^). For mandibles subjected to considerable wear, *W* = 20 *µ*m, the minimum required cutting forces would thus be shifted up by *≈* 40 mN for all leaves. For a 30 mg worker, the fraction of cuttable leaves would be virtually unaffected (99.5 %), whereas a 3 mg worker could cut from almost 50 % of tropical leaves with pristine mandibles to less than 10 % with worn mandibles.

The significant increase of required cutting force with wear, and the conjectured reduction in cuttable plant species, likely necessitates behavioural adaptations, and may partially explain ‘age polyethism’, i.e. systematic changes in task preferences with worker age. Indeed, leaf-cutter ants with worn mandibles cut at significantly lower speeds and are more likely to carry rather than cut [27]; the oldest colony workers may cease foraging altogether, and switch to mechanically less demanding tasks such as waste disposal [50, 94]. The role of wear in determining the mechanical performance of leaf-cutter ants in particular and herbivorous insects in general is worthy of considerably more attention than it has received [44, 53, 55, 57, 60, 95–97].

### Biomechanics of cutting – how sharp are ant mandibles?

The size-invariance of cutting forces and their strong sensitivity to wear have biological consequences. From a mechanical perspective, both results may be surprising at first glance, and thus call for a more thorough evaluation. Intuitively, it appears reasonable to expect that mandibles of larger workers require a larger force to cut the same material. Indeed, the force required to fracture thin or thick model ‘targets’ with biological puncture tools increases significantly with characteristic dimensions of the tool, such as the tip diameter [12, 14]. The expectation that tool size influences mechanical performance is closely tied to the notion of tool ‘sharpness’. However, a robust definition of sharpness as such is not a trivial task, as suitably illustrated by the large number of sharpness metrics suggested in the literature [e. g. 13, 14, 17, 42, 74, 98–101]. In order to rationalise our experimental results qualitatively and quantitatively, we first note that even an arbitrarily sharp mandible will not cut with arbitrarily small force. Cutting is akin to fracture, in the sense that it results in the creation of new surface area. Each unit area of new surface incurs an energy cost *dU*_*A*_, and the work which provides this energy has to be supplied by the externally applied load, so that, from a simple virtual work argument, *dU*_*ext*_ = *dU*_*A*_. Thus, and without loss of generality, the force *F* required to cut a slap of thickness *t* is bound from below by *F∼ G*_*c*_*t*, where *G*_*c*_ is the energy per unit area of new surface, a characteristic material property [41, 68, 71, 102]. For our experiments with **PDMS**, *G*_*c*_ *∼* 100 J m^2^ and *t ∼ 200 µm* (see methods), so that *F ∼* 20 mN. This simple argument lends itself to a definition of an intuitive, quantitative, and functionally relevant index for sharpness, *S*: the required cutting force is equal to the minimum possible force, and independent of tool geometry, if and if only the dimensionless group *S* = *G*_*c*_*tF*^*−*1^ is unity; the cutting tool may then be considered ideally sharp [for a conceptually similar suggestion, see 101]. The fracture forces measured for pristine mandibles of small workers are indeed very close to this theoretical minimum (see Fig. 2B), suggesting that a further reduction in cutting force through changes in mandible morphology may not be possible. Thus, pristine mandibles of small workers appear ideally sharp, *S* ≈1, at least for PDMS (see below for a generalisation of this argument). In contrast, pristine mandibles of larger workers, scalpel blades and the most worn mandibles of foragers have a functional sharpness index *S* between 2/3 and 1/5; in other words, cutting (and fracture) forces are between 50-500% larger than the theoretical minimum, hinting at contributions from cutting tool geometry. The next task is thus to rationalise the putative influence of mandible geometry on cutting force.

The energy associated with the creation of new surface is not the only energy the external force has to supply. Friction, plasticity or sheet bending each carry their own energetic demands, so reducing the fraction of the external work available to drive the cut, *dU*_*ext*_ *− dU*_*l*_ = *dU*_*cut*_ [102]. Some of these costs, for example due to elastic sheet bending or sidewall friction, can be accounted for by drawing the mandible through the cut again, and are thus removed in the fracture force [see Fig. 3B, 11]; but others, related to the direct interaction between the mandible cutting edge and the material close to the crack-tip, likely remain. The simplest possible assumption is that tool geometry can be characterised by a single characteristic length scale, *R* [e. g. 11, 68, 103–105]. From dimensional arguments, this length scale will compete with a characteristic material length scale. In fracture mechanics, the typical length scale is given by the ratio between *G*_*c*_ and a characteristic stress *σ*_*c*_, which may be interpreted physically as a critical crack tip opening displacement, or as the size of a crack process zone in which non-linear mechanisms consume additional energy [e. g. 11, 104, 106–108]. Thus, for this simplest case, dimensional arguments suggest that the additional energy term will be of the form *dU*_*l*_ *∼ Cσ*_*c*_*Rtdx*, where *C* is a dimensionless constant. The fracture force now reads:

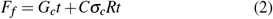

from which the functional sharpness index follows as:

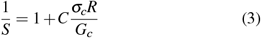

In both equations, the first term represents the unavoidable cost arising from fracture alone; the second term accounts for additional costs linked to tool geometry. For simple geometries such as a cylindrical wire, an exact analysis is possible,and yields *C* = (1 + *µ*), where *µ* is the coefficient of friction [41]. For our experiments, we equate *σ*_*c*_ with the ultimate ten-sile strength of PDMS [about 4 MPa, 70], and assume that the friction coefficient of mandibles on PDMS is similar to values for steel, *µ ≈* 1 [109, 110]. The geometry-dependent term 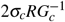 then accounts for half of the cutting force, *S* = 0.5, if the characteristic length is 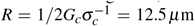. A typical choice for *R* is the radius of the cutting edge [e. g. 11, 12, 14, 68, 98], and indeed, our rather approximate calculation is in remarkable agreement with direct measurements of the cutting edge radius of worn mandibles in *A. cephalotes*,

*R ≈* 17 *µ*m [27]. Pristine mandibles, in turn, may have a cutting edge radius as small as 50 nm [27], so that *S* = 0.996 *≈*1, in seeming agreement with the observation that the pristine mandibles of the smallest workers approach the theoretical minimum cutting force for PDMS (see Fig. 2B). The simple definition of sharpness suggested in Eq. 3 thus has the advantage that it is based on mechanical analysis instead of empirical correlation with observed mechanical performance, that it clearly separates material- and tool-dependent contributions to sharpness, and that its magnitude has a clear physical interpretation.

From this cursory analysis, we may surmise that fracture forces are effectively independent of mandible geometry if 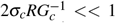, but grow in proportion to *R* ∝ *m*^1*/*3^ for 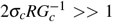 [68, 103]. These limits thus delineate two regimes characterised by geometric invariance and length scaling of cutting forces, respectively, and in practise, the scaling of cutting forces with ***R*** may fall anywhere in between. This result may be put to use in two ways.

First, and in combination with our experimental data, it allows an approximate assessment of the parsimonious but unverified hypothesis that the characteristic mandible dimension ***R*** is isometric, i. e. ***R*** ∝ *m*^1*/*3^. Plausible alternative hypotheses may be derived. For example, the tip radii of insect claws depart from isometry and scale as ***R*** ∝ *m*^1*/*2^, presumably to ensure that tip stresses remain size-invariant [111]. In direct analogy, it is conceivable that pristine mandible cutting edge radii show a scaling shallower than isometry, or are even size-invariant. To test the hypothesis of isometry, we estimate the cutting edge radius *R* from the cutting force measured for a pristine mandible of the largest workers (40 mg in body mass), via *F*_*c*_ = *G*_*c*_*t* + 2*σ*_*c*_*Rt*, yielding *R*_40_ *≈*5 *µ*m. Next, we use this result to extract a proxy for the proportionality constant *a*, invoking the null hypothesis of isometry, *R* = *am*^1*/*3^, and then predict the variation of cutting force across the callow size range from 2.1-46.4 mg, using Eq. 2. An OLS regression on log_10_-transformed predictions yields an intercept of 1.33 and a slope of 0.07, remarkably close to the experimental values of 1.32 and 0.09 (units: mN, mg; see Table 1). Our experimental results are thus consistent with isometry of the mandible cutting edge radius. Although *R* may vary by as much as a factor of 30^0.33^ ≈3 across the size range investigated in this study, cutting forces vary only little with size, because even large mandibles satisfy 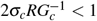. However, the considerable variation in our data even for pristine mandibles limits the statistical power to establishing consistency, and direct experimental assessment, for example via scanning electron microscopy [12, 27], is necessary to firmly establish isometry.

Second, Eq. 2 can be put to work to assess whether the size-invariance of cutting forces observed for a synthetic material such as PDMS may extend to natural materials typically cut by leaf-cutter ants. To this end, we extract proxies for the median *G*_*c*_ *≈* 400 N m^-1^, *t* = 200 *µ*m and *σ*_*c*_ ≈ 3 N mm^-2^ from an extensive study on the leaf lamina of about 1000 tropical plant species [8], and again use Eq. 2 to predict the expected scaling of cutting forces. We find an intercept of 1.9 and a slope of 0.02. Thus, the size-dependence of the net cutting force in natural materials may be even weaker than for PDMS, because leaves have a higher toughness, but similar ultimate strength, so that 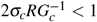, and the geometry-independent term in Eq. 2 dominates. We stress that this analysis is approximate, and cutting of plant leaves may for example incur larger bending costs, because they are much stiffer. Preliminary support is however available from cutting force measure-ments with laurel leaves. Based on the median tropical leaf with *G*_*c*_ = 400 N m^-1^, and *σ*_*c*_ = 3 N mm^-2^, and the lamina thickness of laurel, *t ≈*250 *µ*m, Eq. 2 predicts cutting forces for a pristine mandible with *R* = 5 *µ*m and a worn mandible with *R* = 12.5 *µ*m of 108 and 119 mN, respectively, in reasonable agreement with our experimental results (see Fig. 3A). Thus, a difference in cutting edge radius that would increase cutting forces in PDMS by about 40% increases those for the median leaf by a mere 10%. Although the simple model based on dimensional arguments appears to quantitatively capture salient features of our experimental data, more thorough experimental validation, including cutting measurements with a range of natural materials and direct measurements of cutting edge radii, are in order.

The putatively weak size-dependence of mandible cutting forces for natural materials has two consequences worthy of brief discussion. First, it implies that mandible wear needs to be more severe in order to have an appreciable effect on cutting forces. As an illustrative example, *S* = 0.5, corresponding to a doubling of the required cutting force, occurs for *R ≈*12.5 *µ*m in PDMS; the equivalent radius for the median tropical leaf is

*R≈* 67 *µ*m – about five times larger. However, there is robust evidence that wear affects leaf-cutter ant performance even when cutting natural materials: the average fracture force required for forager mandibles to cut laurel leaf lamina was about 30 mN higher than for callow mandibles (see Fig. 3A), and similar results were reported by Schofield et al. [27] for *A. cephalotes* workers and *Prunus lusitanica* leaves; leaf-cutter ants with worn mandibles cut at significantly lower speeds [27, see also 53 for similar results on leaf beetles]; and leaf-cutter ants with worn mandibles show changes in task preferences [27, 50, 94]. Clearly, the role of wear in modulating cutting forces of natural materials requires further experimental investigation. Second, and conversely, it suggests that even moderately small cutting edge radii may suffice to achieve *S* ≈1 For example, for *R* = 1 *µ*m, *S* = 0.99*≈* 1, and even for *R* = 10 *µ*m, *S* = 0.87, still within 15% of the maximum sharpness for cutting the median leaf. Thus, selection pressure on materials and edge geometry for the cutting tools of small animals may be less strong than previously suggested [12, 47].

### Conclusions and outlook

The ability to cut leaves involves complex interactions between worker size, bite force capacity, wear-dependent cutting forces, plant-material properties and adaptive cutting behaviour. We tried to untangle this complexity, by removing the confounding effects of material heterogeneity and non-linear mandible motion, and studied the effects of worker size across two experimental groups with varying levels of mandibular wear.

Although smaller ants may experience a larger increase in cutting force from pristine to worn mandibles, cutting forces were still largely size-independent, in contrast to our initial hypothesis. The ability to cut leaves is thus mostly affected by size-dependent bite forces, plant-material properties, and mandibular wear, so that larger ants require a substantially smaller fraction of their maximum bite force to cut the same material. In agreement with our second hypothesis, the effects of wear on cutting force can be substantial, which may strongly reduce the range of accessible plant tissues for small workers.

Pristine mandibles of callow workers are exceedingly ‘sharp’, and even mandibles with moderate levels of mandibular wear require similar forces to the ‘sharpest’ pristine scalpel blade; these results indicate morphological adaptations of leaf-cutter ant mandibles to the high mechanical demands of cutting [27, 45].

A natural extension to this work would be to use other materials as cutting substrate, and to test quantitative predictions on cutting force variation and cutting edge geometry for a broader selection of biologically relevant substrates. A careful inspection of the mandibular cutting blade, in combination with mandible abrasion experiments, could yield important insights into the mechanisms of wear resistance in insects [43, 45].

We hope that the findings of this study will help to increase our understanding of size-specific foraging preferences in leafcutter ants, and more generally, may provide a framework to discuss the relative importance of tool geometry vs material properties in biological cutting.

## Supporting information

Video: Leaf-cutter ant cutting bramble

## Acknowledgments

We thank Flavio Roces, who for kindly provided the ant colonies used in this study, and Franka Nauert, Aurèlie Levillain and Andrea Attipoe for their preliminary work on the fibre-optic setup. This study is part of a project that has received funding from the European Research Council (ERC) under the European Union’s Horizon 2020 research and innovation programme (Grant agreement No. 851705) awarded to DL.

