## Supplementary Information for "Biomechanics of cutting: sharpness, wear sensitivity, and the scaling of cutting forces in leaf-cutter ant mandibles"

### Supplementary Materials

### Tooth length measurements and wear determination

In order to estimate mandibular wear, we extracted the lengths of the most distal tooth, $T_1$, and second most distal tooth, $T_2$, of the mandible blade from photographic images, following Schofield et al. [see 1, and Fig. 1 in the main text]. Tooth length was measured perpendicular to the line connecting the most distal tooth gap, also called 'V-blade', and the most proximal tooth gap; the length of this line determined the mandible blade length, $L_{mb}$. To establish the relationship between $L_{mb}$, $T_1$, and $T_2$ with body mass, we performed OLS regressions on $\log_{10}$-transformed values (see Table 1 and Fig. 2B). Based on the regression results obtained for the pristine teeth of callows, we calculated tooth length differences and mandibular wear (see Eq. 1 in the main text). Notably, all length parameters were close to isometric scaling, $m^{0.33}$, apart from $T_2$ for pristine callow mandibles, which was proportional to $m^{0.12}$ (see Table 1); this result suggests changes in mandible blade curvature across sizes [2, and N Imirzian, F Püffel and D Labonte, in preparation, 3].

[3] Imirzian N, Püffel F, Roces F, Labonte D. Large deformation diffeomorphic metric mapping reveals 3d shape variation of head capsules and mandibles in *Atta vollenweideri* leaf-cutter ants. Manuscript in preparation.

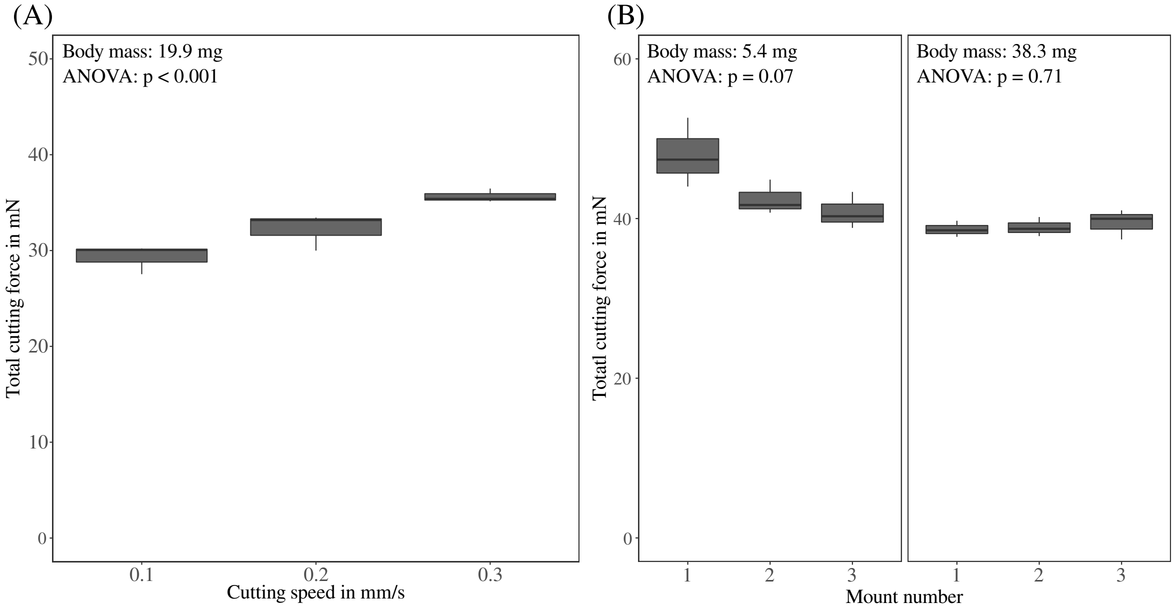


Figure 1 | **(A)** To quantify the effect of cutting speed on mandibular cutting forces, the mandible of a single forager (body mass 19.9 mg) was tested at 0.1, 0.2 and 0.3 mm $s^{-1}$ for three repetitions each, approximately covering the range of cutting speeds observed in foraging *Atta*. The total cutting forces increased significantly with speed from a mean of 29 to 36 mN (or about 20 %; see main text for statistics). Although significant, this increase is small compared to the overall variation across callow and forager mandibles (19 - 172 mN), and thus perhaps of limited biological significance. **(B)** Similarly, the variation from mounting was small in comparison to inter-individual variation, as indicated by the coefficient of variation, $CV$: $CV_{\text{small}} = 0.10$, $CV_{\text{large}} = 0.03$, $CV_{\text{foragers}} = 0.52$. Notably, the $CV$ was larger for the smaller ant, suggesting that consistent mandible alignment was easier for larger mandibles.

Table 1 | Results of Ordinary Least Squares regressions describing the relationship between various mandible dimensions in $\mu$m and body mass in mg. All regressions were performed on $\log_{10}$-transformed data; 95 % confidence intervals are shown in parentheses.

| Quantity | Group | Elevation | Slope | $R^2$ |
|---|---|---|---|---|
| $L_{mb}$ | Foragers | 2.63 (2.62, 2.65) | 0.31 (0.30, 0.33) | 0.99 |
| $L_{mb}$ | Callows | 2.66 (2.64, 2.68) | 0.29 (0.27, 0.31) | 0.96 |
| $T_1$ | Foragers | 1.95 (1.76, 2.14) | 0.38 (0.20, 0.56) | 0.47 |
| $T_1$ | Callows | 2.12 (2.08, 2.15) | 0.32 (0.29, 0.35) | 0.92 |
| $T_2$ | Foragers | 1.50 (1.32, 1.68) | 0.25 (0.08, 0.42) | 0.30 |
| $T_2$ | Callows | 1.70 (1.64, 1.77) | 0.12 (0.06, 0.18) | 0.28 |

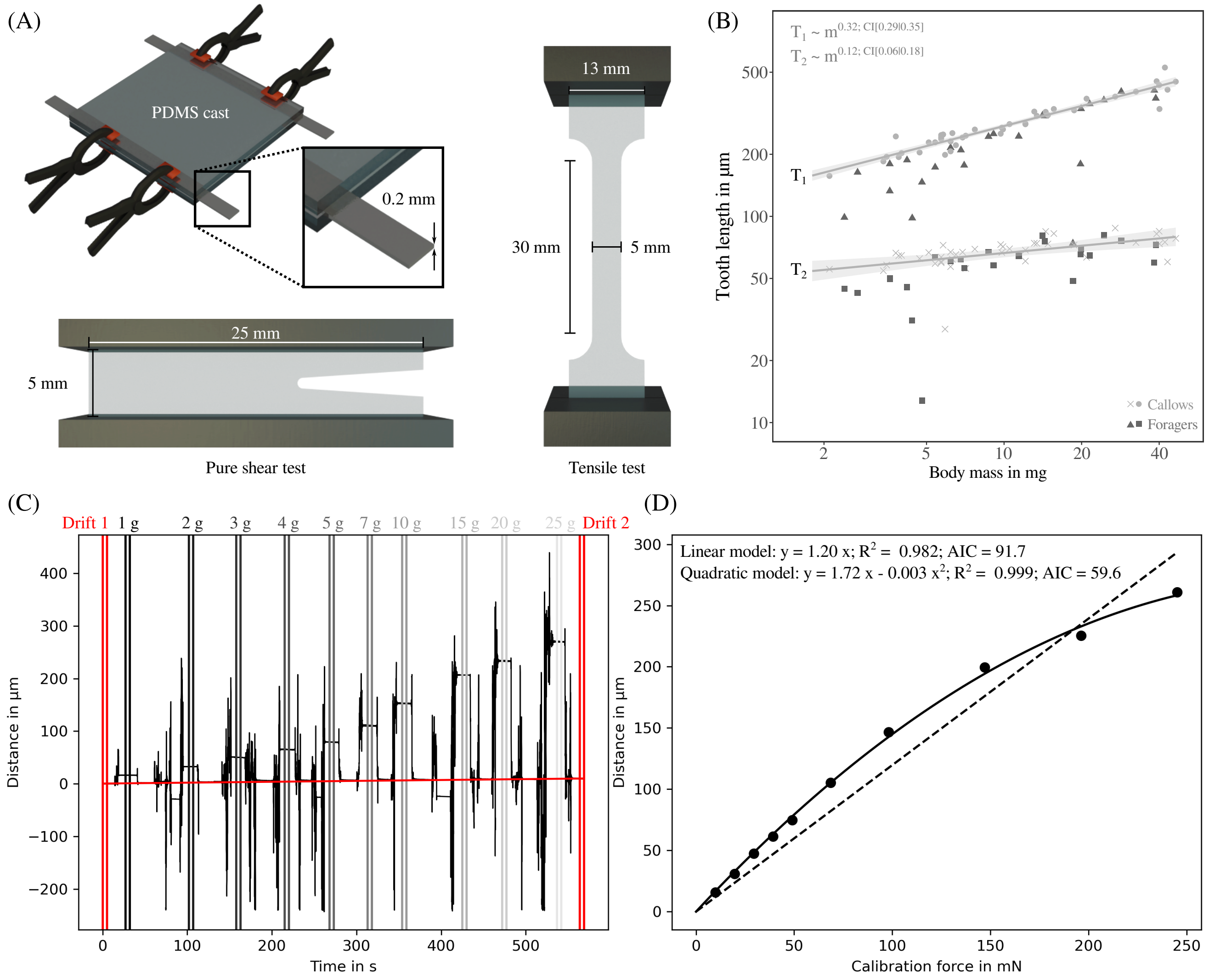


Figure 2 | **(A)** For the cutting experiments, we used PDMS sheets as controlled test substrate. Individual sheets were formed between two glass plates, oven-cured, and mechanically characterised via pure shear and tensile tests (see main text for more details). **(B)** In order to quantify mandibular wear, we calculated a wear index based on absolute length changes to the two most distal teeth, $T_1$ and $T_2$, respectively [1]. The pristine length of the most distal tooth increased as expected from isometry; the pristine length of the second most distal tooth increased with negative allometry (see Table 1). **(C)** To calibrate the force sensor used for cutting experiments, we suspended a series of calibration weights from the free end of the bending beam (see methods). From the distance-time curve, we extracted the average distance across 5 s for each weight, and implemented a linear drift correction (red line). **(D)** We tested different regression models to characterise the relationship between distance and calibration force: the quadratic model yielded a lower Akaike Information Criterion and was hence preferred over the linear model.

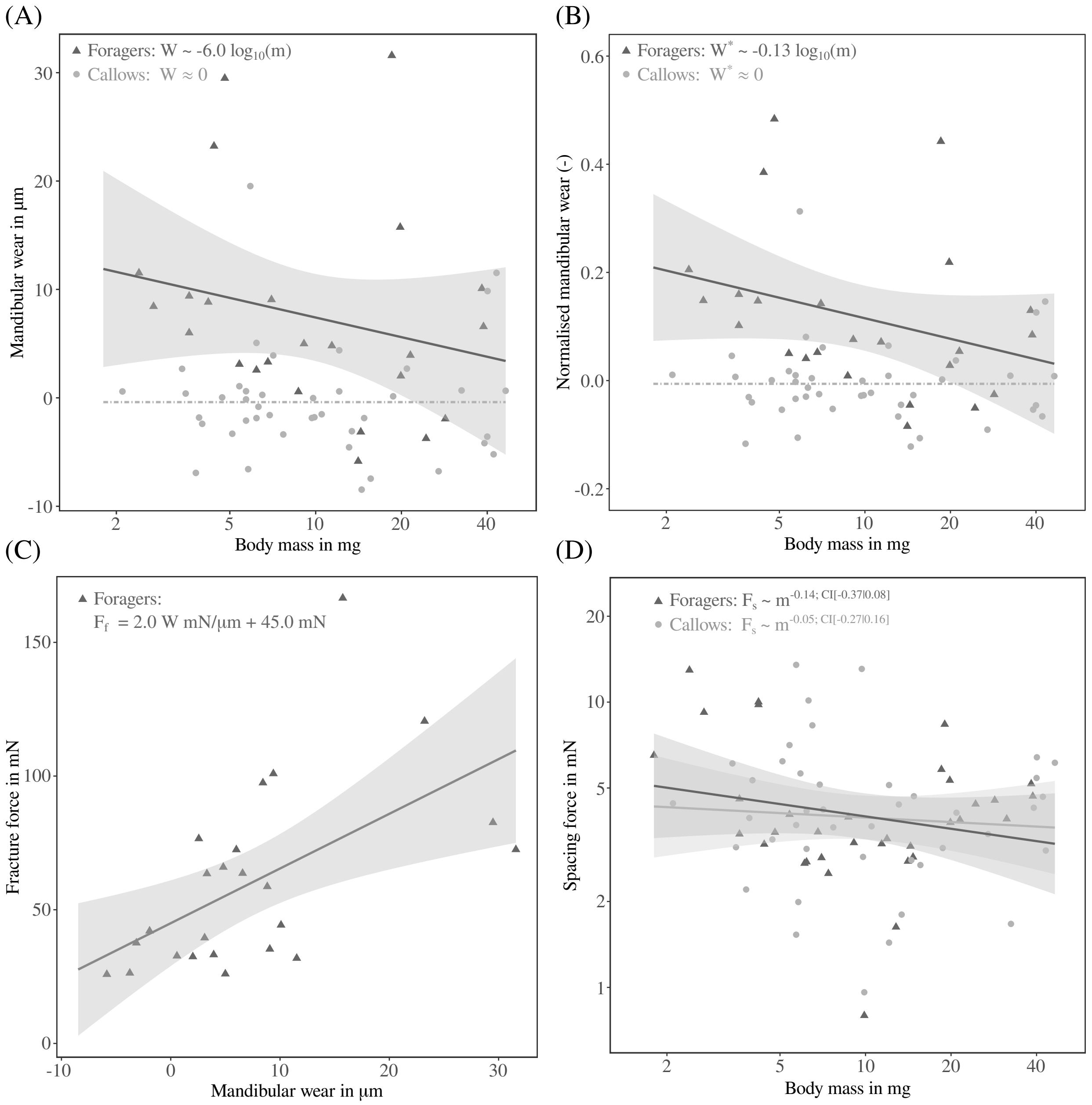


Figure 3 | **(A)** We calculated a simple mandibular wear index, $W$, defined as the weighted average length change of the two most distal teeth [see 1, and Eq. 1 in the main text]. For foragers (n = 24), the average absolute wear index was $8 \pm 10\ \mu$m, independent of body mass; for callows (n = 42), $W \approx 0$, by definition (units: $\mu$m, mg). **(B)** Similarly, the relative wear index for foragers, $W^{\star}$, normalised with the pristine length of the second most distal tooth, was independent of body mass, but tended to be larger for smaller workers. **(C)** In analogy with Fig. 3C in the main text, the fracture force, $F_f$, increased significantly with wear (n = 24, Ordinary Least Squares regression on forager data: slope = 2.05, 95 % CI (0.71 | 3.38), $R^2$ = 0.31), and at a similar rate as the total cutting force. **(D)** Spacing forces were generally small for mandible-PDMS cutting experiments, with neither significant differences between experimental groups, nor significant size-effects (n = 76).
